# Impaired complement regulation drives chronic lung allograft dysfunction after lung transplantation

**DOI:** 10.1101/2024.11.17.623951

**Authors:** Hrishikesh S. Kulkarni, Laneshia K. Tague, Daniel R. Calabrese, Fuyi Liao, Zhiyi Liu, Lorena Garnica, Nishanth Shankar, Xiaobo Wu, Devesha H. Kulkarni, Cory Bernardt, Derek Byers, Catherine Chen, Howard J. Huang, Chad A. Witt, Ramsey R. Hachem, Daniel Kreisel, John P. Atkinson, John R. Greenland, Andrew E. Gelman

**Author notes:** **Corresponding Authors:** Hrishikesh Kulkarni, MD, MSCI Division of Pulmonary, Critical Care and Sleep Medicine University of California, Los Angeles, 10833 Le Conte Ave, 52-257 CHS, Mail code: 169017 Los Angeles, CA 90095, Andrew E. Gelman, PhD Mary Culver Department of Surgery Washington University School of Medicine 660 S Euclid Ave St. Louis, MO 63110.

## Abstract

A greater understanding of chronic lung allograft dysfunction (CLAD) pathobiology, the primary cause of mortality after lung transplantation, is needed to improve outcomes. The complement system links innate to adaptive immune responses and is activated early post-lung transplantation to form the C3 convertase, a critical enzyme that cleaves the central complement component C3. We hypothesized that LTx recipients with a genetic predisposition to enhanced complement activation have worse CLAD-free survival mediated through increased adaptive alloimmunity. We interrogated a known functional C3 polymorphism (C3R102G) that increases complement activation through impaired C3 convertase inactivation in two independent LTx recipient cohorts. C3R102G, identified in at least one out of three LTx recipients, was associated with worse CLAD-free survival, particularly in the subset of recipients who developed donor-specific antibodies (DSA). In a mouse orthotopic lung transplantation model, impaired recipient complement regulation resulted in more severe obstructive airway lesions when compared to wildtype controls, despite only moderate differences in graft-infiltrating effector T cells. Impaired complement regulation promoted the intragraft accumulation of memory B cells and antibody-secreting cells, resulting in increased DSA levels. In summary, genetic predisposition to complement activation is associated with B cell activation and worse CLAD-free survival.

**BRIEF SUMMARY:** Lung transplant recipients genetically predisposed to impaired complement regulation demonstrate worse chronic rejection-free survival. This phenotype is associated with intragraft B cell-activation and donor-specific antibodies.

## INTRODUCTION

Chronic lung allograft dysfunction (CLAD) is the primary cause of long-term morbidity and mortality after lung transplantation (1, 2). CLAD commonly presents with a constrictive bronchiolitis pathology and is associated with poor survival (3). However, it is unclear why some lung transplant recipients progress to CLAD whereas others do not. CLAD is also a heterogenous syndrome with greater or lesser involvement of donor-specific antibodies (DSA) and distal parenchyma, leading to antibody mediated rejection (AMR) and restrictive allograft syndrome (RAS) phenotypes (4–6). Cross-talk between innate and adaptive immunity via the complement system may be necessary for driving CLAD (7–9). While T cells are required for acute and chronic rejection, there is increasing evidence that B cells contribute to poor lung transplant survival, potentially through AMR (10–12). Additionally, mouse models of lung transplantation have demonstrated B cell-dependent CLAD-like pathology (13, 14).

The complement system comprises over 50 proteins that are activated, amplified, and deposited onto pathogens and dying cells to facilitate their clearance by phagocytosis (15).The system can also be activated in transplantation via antibody-dependent and antibody-independent pathways (16–19). C3a and C5a are cleaved from C3 and C5 upon activating this cascade and can engage with their cognate receptors to influence alloimmune responses (20). Engagement of these receptors also drives profibrotic responses (21, 22). At the same time, the key complement protein C3 is required for the survival of both immune (23) and non-immune (24–26) cells, and modulates effector cell function (27).

A series of both membrane-bound and fluid-phase regulators control complement activation and facilitate tissue homeostasis (28). Genetic and acquired deficiencies in these regulators increase complement activation, resulting in augmented tissue damage (29, 30). CD46, a membrane regulator, serves as a cofactor for fluid-phase regulators such as Factor I to cleave C3b and C4b, and inactivates the C3 convertase to attenuate complement activation. Crry, a murine ortholog of CD46, has been shown to prevent complement activation on the vascular endothelium (31). Crry is downregulated during lung injury (32), and has also been used to ameliorate lung injury (33, 34).

The role of complement regulation in CLAD remains undefined. Given the centrality of C3 in the cascade, we analyzed a functional polymorphism in C3 (*rs2231099*, C3 R102G) that has a known minor allele frequency of at least 10% in several large cohorts. This polymorphism results in increased complement activation due to the inability of the fluid-phase regulator Factor H to inactivate the C3 convertase (35). Previous work has shown that the C3 R102G polymorphism is linked to the development of age-related macular degeneration and other complement-related conditions (36). We hypothesized that C3 R102G is associated with CLAD development. We show that recipients carrying the C3 R102G polymorphism in two independent human cohorts have worse CLAD-free survival, and these outcomes are more pronounced in the setting of DSA. Additionally, using a mouse orthotopic lung transplantation model, we demonstrate that recipients deficient in the complement regulator Crry develop worse signs of obliterative airway disease that are linked to elevated intragraft B cell activation and high DSA levels.

## RESULTS

### A functional polymorphism in complement component C3 gene is observed in at least 1 out of 3 lung transplant recipients

Within the lung transplantation cohort at Washington University/Barnes-Jewish Hospital (BJH), we identified 392 participants with available genotyping (**Figure 1A**). Among these recipients, 245 (62.5%) had wild-type C3 (G/G); thus, the recipient *rs2230199* (G>C) SNP had a minor allele frequency (allele C) of 37.5%. Baseline characteristics for participants stratified by genotype are shown in **Table 1** and were similar across both genotypes.

**Figure 1.**
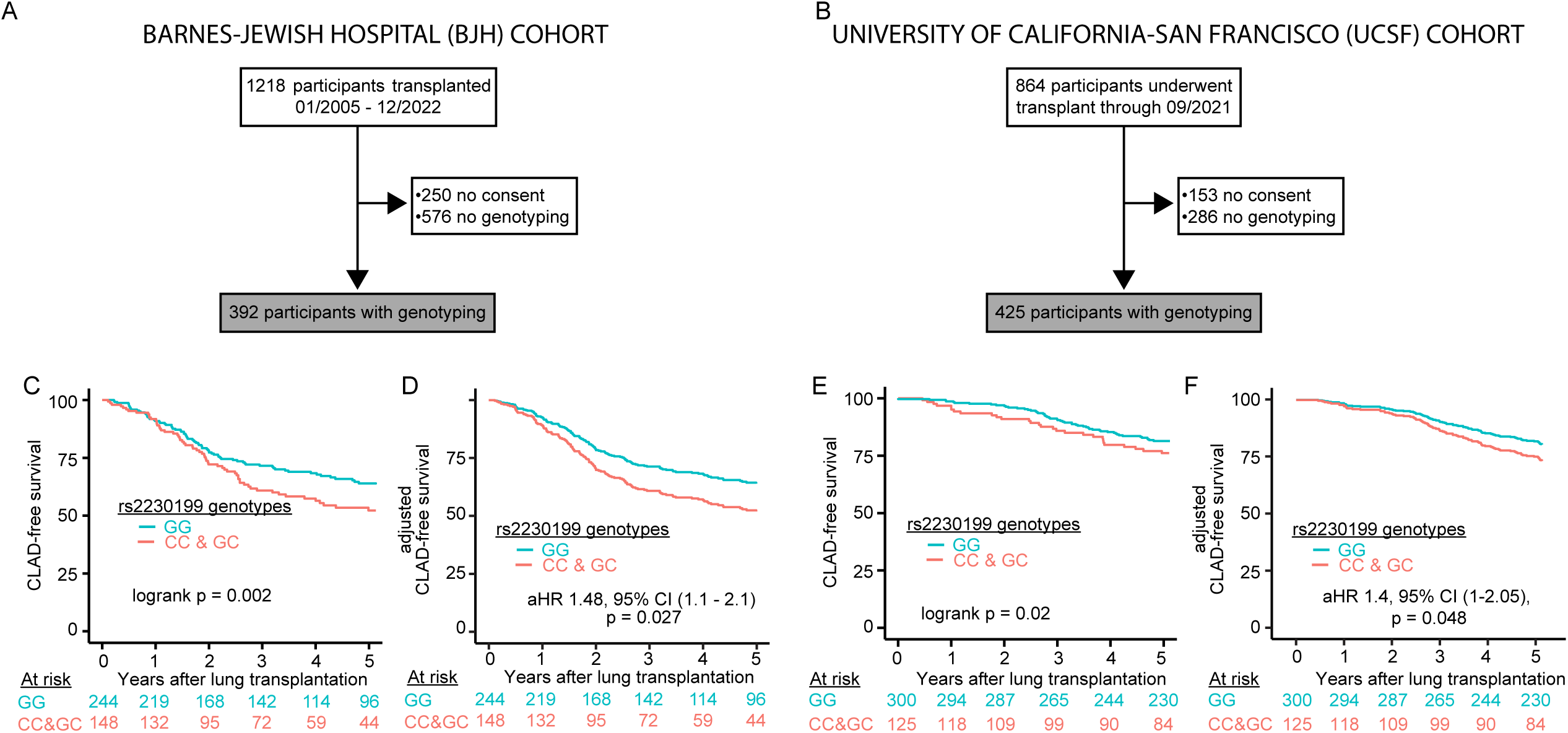
A functional C3 polymorphism confers increased risk of chronic lung allograft dysfunction (CLAD) or death in two independent cohorts. CONSORT diagrams for (A) Washington University/Barnes-Jewish Hospital (BJH) and (B) University of California-San Francisco (UCSF) cohorts. (C and E) Kaplan Meier plot of survival from CLAD or death in the BJH (C) and UCSF (E) cohort stratified by *rs2230199* genotypes. (D and F) Kaplan Meier plot based on the multivariate Cox Regression analysis of CLAD-free Survival in the BJH (D) and UCSF (F) cohorts by *rs2230199* status. P value represents log-rank test. aHR: adjusted hazard ratio; CI: confidence intervals.

**Table 1.**
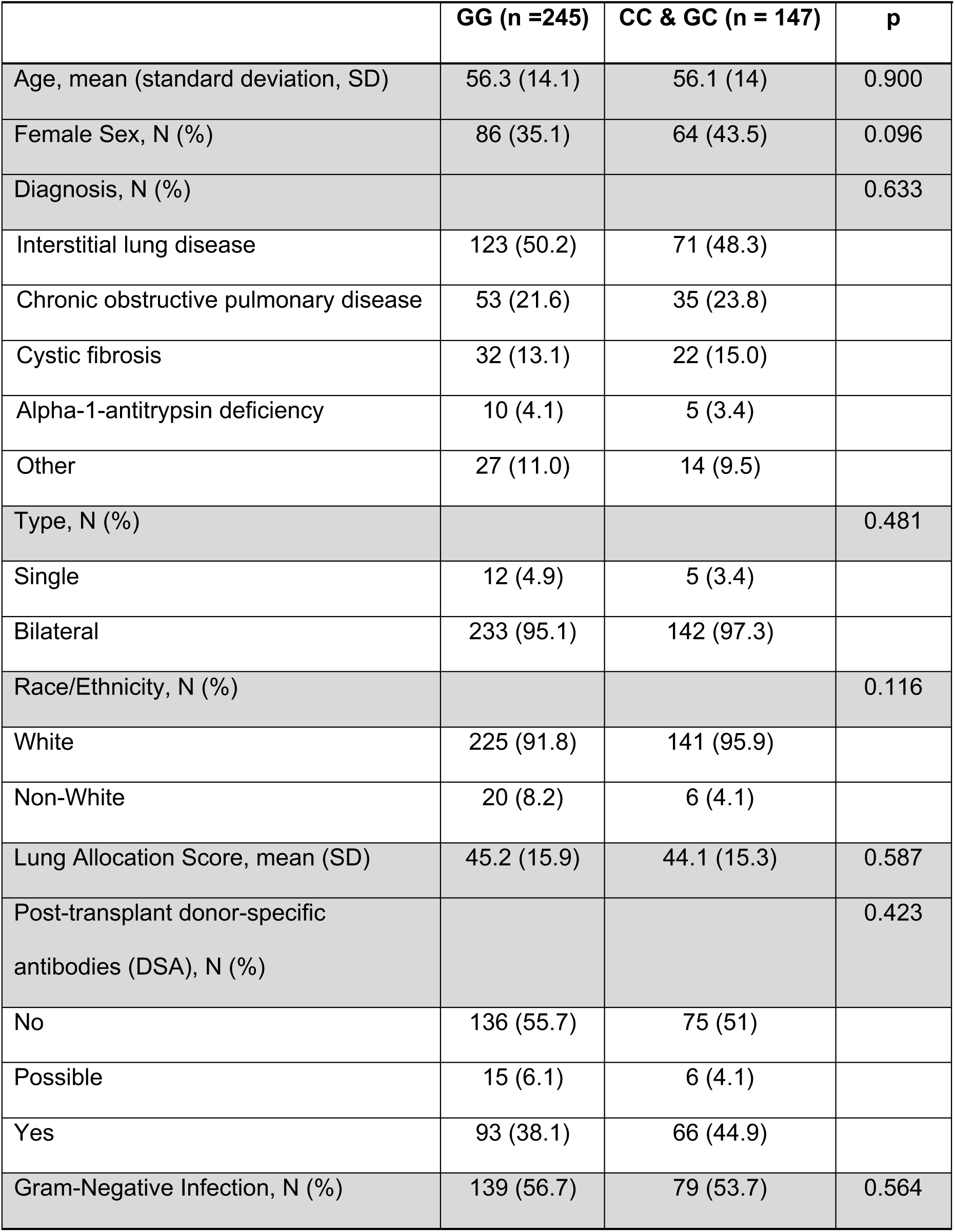
Barnes-Jewish Hospital (BJH) cohort baseline characteristics.

Within the UCSF lung transplantation cohort, we identified 425 participants with available genotyping (**Figure 1B**). The recipient *rs2230199* (G>C) SNP had a minor allele frequency (allele C) of 29.4%. Baseline characteristics for participants stratified by genotype are shown in **Table 2**. Notably, we found higher minor allele frequencies among Caucasian recipients and lower minor allele frequencies among African American recipients. These distributions are consistent with those reported in aggregated global genomic databases (37). There were no other differences in lung transplant baseline characteristics across genotypes.

**Table 2.** UCSF cohort baseline characteristics.

|  | **GG (n=300)** | **CC & GC (n = 125)** | **p** |
| --- | --- | --- | --- |
| Age, mean (standard deviation, SD) | 54.7 (12) | 56 (11.9) | 0.34 |
| Female Sex, N (%) | 130 (43.3) | 58 (46.4) | 0.67 |
| Diagnosis, N (%) |  |  | 0.684 |
| Interstitial lung disease | 203 (67.7) | 85 (68) |  |
| Chronic obstructive pulmonary disease | 53 (17.7) | 22 (17.6) |  |
| Cystic fibrosis | 22 (7.3) | 13 (10.4) |  |
| Alpha-1-antitrypsin deficiency | 7 (2.3) | 1 (0.8) |  |
| Pulmonary hypertension | 14 (4.7) | 4 (3.2) |  |
| Type, N (%) |  |  | 0.154 |
| Single | 31 (10.3) | 6 (4.8) |  |
| Bilateral | 257 (85.7) | 112 (89.6) |  |
| Heart-Lung | 12 (4) | 7 (5.6) |  |
| Race/Ethnicity, N (%) |  |  | 0.007 |
| White | 198 (66) | 101 (80.8) |  |
| Non-White | 102 (44) | 24 (119.2) |  |
| Lung Allocation Score, mean (SD) | 45.4 (17.5) | 44.1 (18.6) | 0.49 |
| Post-transplant donor specific antibodies (DSA), N (%) |  |  | 0.18 |
| No | 233 (77.7) | 105 (84) |  |
| Yes | 67 (22.3) | 20 (16) |  |
| Gram-Negative Infection, N (%) | 113 (37.7) | 41 (32.8) | 0.4 |

### C3 R102G confers increased risk of CLAD or death in two independent cohorts

We next examined whether the *rs2230199* C3 minor allele, which predisposes to increased complement activation, was associated with decreased CLAD-free survival. In the BJH cohort, the C3 R102G polymorphism was associated with an increased risk for CLAD or death (**Figure 1C**). In a multivariable Cox-proportional hazards model consisting of C3 *rs2230199* genotype, age, sex, race, transplant diagnosis, DSA and Gram-negative infection, C3 R102G remained associated with increased risk of CLAD or death (**Figure 1D**, **Table 3**). The presence of post-transplant DSA (1.74 aHR, 1.24 – 2.45 95% CI, p=0.001) or post-transplant Gram-negative infection (1.44 aHR, 1.01 – 2.06 95% CI, p=0.044) were also associated with a significantly increased risk of CLAD or death. A sensitivity analysis removing the presence of DSA among the list of variables demonstrated that C3 R102G was still associated with an increased risk of CLAD or death (1.52 aHR, 1.08 – 2.15 95% CI, p=0.016). A similar analysis removing the presence of Gram-negative infection among the list of variables demonstrated that C3 R102G remained associated with an increased risk of CLAD or death (1.43 aHR, 1.02 – 2.02 95% CI, p=0.04).

**Table 3.** BJH Multivariate Cox Regression Analysis of CLAD-free Survival by *rs2230199* status (n=390)

| Variable | aHR (95% CI) | p |
| --- | --- | --- |
| Age | 1.00 (0.99-1.02) | 0.628 |
| Female Sex | 1.32 (0.92-1.89) | 0.138 |
| Non-white Race | 1.46 (0.76-2.82) | 0.255 |
| Transplant Diagnosis |  | 0.868 |
| Interstitial lung disease (reference) | - |  |
| Chronic obstructive pulmonary disease | 1.09 (0.71 – 1.66) |  |
| Cystic fibrosis | 0.84 (0.41 – 1.69) |  |
| Alpha-1-antitrypsin deficiency | 0.59 (0.18 – 1.90) |  |
| Other | 1.04 (0.92 – 1.89) |  |
| Post-transplant donor specific antibodies |  | 0.001 |
| Absent (reference) | - |  |
| Present | 1.74 (1.24-2.45) |  |
| Post-transplant Gram-negative infection |  | 0.044 |
| Absent (reference) | - |  |
| Present | 1.44 (1.01-2.06) |  |
| C3 *rs2230199* genotype |  | 0.027 |
| GG (reference) | - |  |
| GC/CC | 1.48 (1.05-2.08) |  |

We examined if R102G is also associated with differential CLAD-free survival in the UCSF cohort. **Figure 1E** shows that participants homozygous for C3 R102G had an increased risk for CLAD or death (1.43 HR, 1 – 2.05 95% CI, adjusted p = 0.048). This multivariable Cox-proportional hazards model also consisted of C3 *rs2230199* genotype, age, sex, race, transplant diagnosis, DSA and Gram-negative infection (**Figure 1F**, **Table 4**).

**Table 4.** UCSF Multivariate Cox Regression Analysis of CLAD-free Survival by *rs2230199* status (n=425)

| Variable | aHR (95% CI) | p |
| --- | --- | --- |
| Age | 1 (0.99-1.02) | 0.337 |
| Female Sex | 1.31 (0.94-1.83) | 0.107 |
| Non-white Race | 0.79 (0.53 – 1.18) | 0.254 |
| Transplant Diagnosis |  | 0.462 |
| ILD (reference) | - |  |
| COPD | 1.33 (0.88 – 2.02) |  |
| CF | 0.68 (0.31 – 1.47) |  |
| Alpha-1-antitrypsin deficiency | 1.2 (0.37 – 3.86) |  |
| Other | 1.01 (0.38 -2.64) |  |
| Post-transplant donor specific antibodies |  | 0.018 |
| Absent (reference) | - |  |
| Present | 1.62 (1.1-2.41) |  |
| Post-transplant Gram-negative infection |  | 0.074 |
| Absent (reference) | - |  |
| Present | 1.4 (0.96 – 1.96) |  |
| C3 *rs2230199* genotype |  | 0.048 |
| GG (reference) | - |  |
| GC/CC | 1.43 (1 – 2.05) |  |

### CLAD-free survival association with recipient C3 R102G is linked to donor specific antibodies (DSA)

Based on work published by us and others that anti-HLA DSA are a known risk factor for CLAD (11, 38), we hypothesized that the increased risk of CLAD or death in recipients with C3 R102G would be limited to those with DSA. In the BJH cohort, 181 (46.2%) lung transplant recipients had definite DSA during their follow-up. CLAD-free survival analysis accounting for both DSA and *rs2230199* genotype demonstrated that recipients with C3 R102G and DSA had the worst CLAD-free survival **(Figure 2A).** Among recipients without DSA, we did not observe a difference in risk for CLAD or death based on the C3 *rs2230199* genotype (1.21 HR, 0.72 – 2.04 95% CI, p = 0.48). However, among the participants who developed DSA and had the C3 R102G polymorphism, there was an increased risk for CLAD or death (1.52 HR, 0.99-2.34 95%CI, p = 0.05), with the difference in survival primarily being observed starting at approximately 2.5 years post-transplantation.

**Figure 2.**
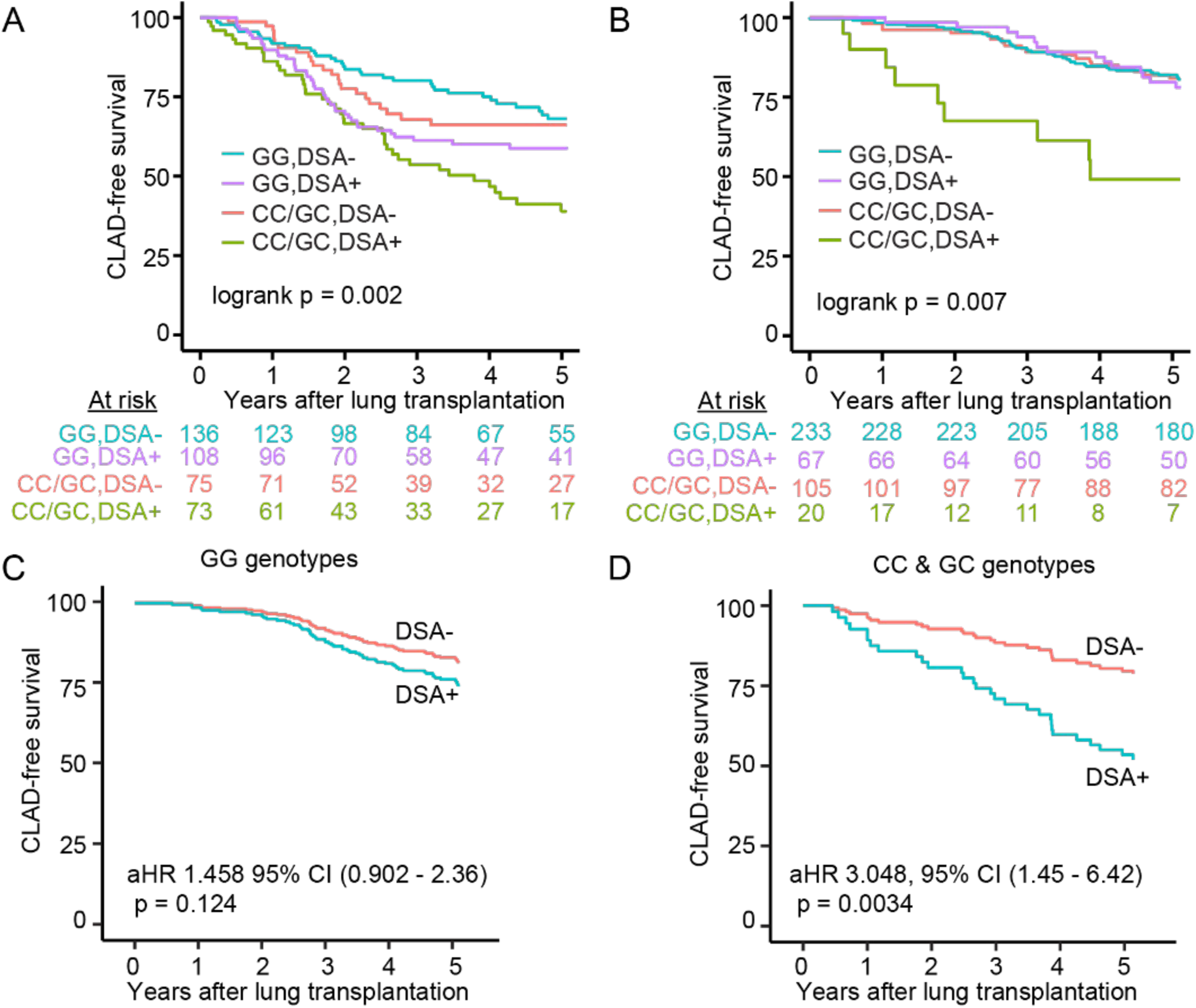
Complement-mediated CLAD or death is dependent on donor-specific antibodies (DSA). CLAD-free survival was worse in the recipients with donor-specific antibodies (DSA) and the C3 R102G polymorphism (CC/GC). The BJH (A) and UCSF (B) cohorts were stratified by DSA-negative and DSA-positive recipients. p value represents log-rank test. (C and D) Kaplan Meier plot based on the multivariate Cox Regression analysis of CLAD-free survival in the UCSF cohort separated by genotype, stratified by DSA status. aHR: adjusted hazard ratio, CI: confidence interval.

These observations also held true in the UCSF cohort, wherein 87 (20.5%) lung transplant recipients had evidence of DSA at some point during their post-transplant course. Among participants without DSA there was no different risk for CLAD or death by the *rs2230199* genotype (**Figure 2B**, 1.2 aHR, 0.85 – 1.9 95% CI, adjusted p = 0.24). However, among the participants who developed DSA and had either CC/GC alleles, there was a 3.2-fold increased risk for CLAD or death (3.2 aHR, 1.6 – 6.2 95% CI, adjusted p = 0.0007). These models were adjusted for age, sex, race, transplant diagnosis, and Gram-negative infection. These findings suggest that the impact of the *rs2230199* SNP is dependent upon DSA development.

We next examined CLAD-free survival separately in recipients based on the presence or absence of C3 R102G. Recipients from the UCSF cohort who lacked C3 R102G (i.e., GG genotype) did not demonstrate a significant difference in CLAD-free survival irrespective of their DSA status (**Figure 2C**). In comparison, recipients with post-transplant DSA and the C risk allele (i.e., CG or CC genotype) were at a threefold higher risk for worse CLAD-free survival after adjusting for age, sex, race, transplant diagnosis, and Gram-negative infection (**Figure 2D**). The results also held true in the BJH cohort, where recipients with post-transplant DSA and the C risk allele demonstrated worse CLAD-free survival (1.9 aHR, 1.1 – 3.3 95% CI, adjusted p = 0.02). These data suggest that the worst long-term outcomes occurred in those harboring the *rs2230199* SNP, who developed post-transplant DSA.

### Impaired complement regulation promotes CLAD in the mouse orthotopic lung transplant model

The C3 R102G polymorphism lacks a synonymous variant in mice. However, mice deficient in Crry (*Crry^-/-^*) have increased complement activation due to impaired Factor I-mediated cleavage of C3b and C4b, resulting in dysregulated C3 convertase formation on the membrane (39). As a result, these mice have more C3 consumption and lower circulating levels of C3 (40, 41), as well as attenuated levels of C3 in their bronchoalveolar lavage (BAL) (**Figure 3A**). Given our prior findings that C3 is required for epithelial cell survival (26) and that C3 R102G predisposes to impaired complement regulation (35), we asked if CLAD development is regulated by complement activation. For this purpose, we utilized a previously established mouse orthotopic lung transplant model of CLAD (42, 43) (**Figure 3B**). Wildtype *Crry^+/+^* and *Crry^-/-^* (Crry-deficient) recipients, both on a B6 background (H-2^b^), were engrafted with major-mismatched left lungs encoding three transgenes (3T-FVB; H-2^q^); a reverse tetracycline activator gene driven by the club cell secretory protein promoter, a Cre recombinase gene under the control of the reverse tetracycline activator and a lox-P activated diphtheria toxin A gene. These transgenes induce graft-specific club cell injury and depletion in response to doxycycline ingestion, potentiating CLAD development in response to alloimmune stress (43).

**Figure 3:**
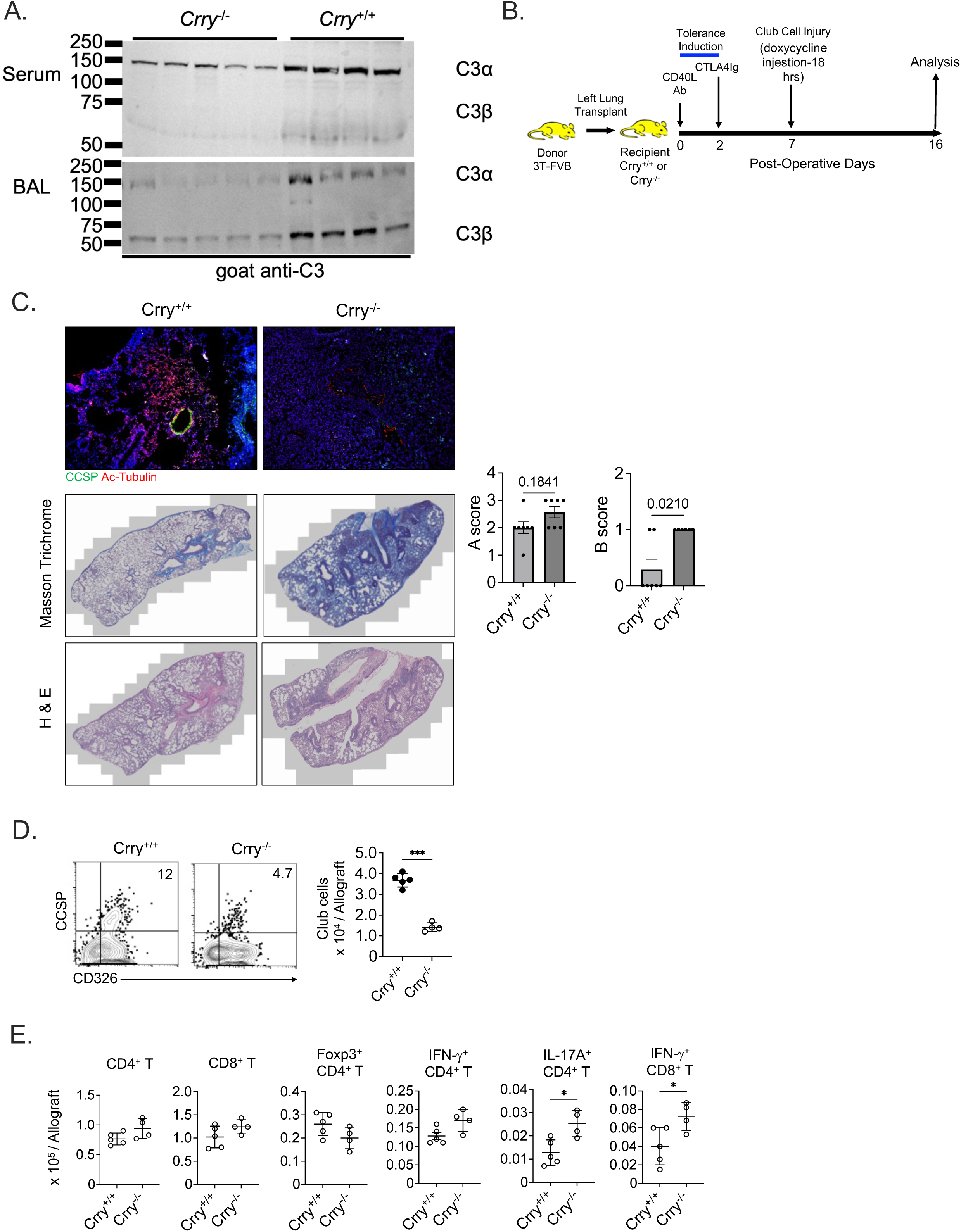
Dysregulated complement activation promotes CLAD in a mouse lung transplantation model. (A) Immunoblot of serum and bronchoalveolar lavage (BAL) C3-alpha and -beta fragments from resting *Crry*^+/+^ and *Crry*^-/-^ mice (N=4/group). The data shown is a representative result of 2 experiments. (B) Mouse orthotopic left lung transplant model of CLAD. (C, upper panel) POD 16 immunofluorescent staining of club and ciliated cells using antibodies specific for club cell secretory protein (CCSP) and acetylated tubulin (Ac-Tubulin), respectively (N=5/group). (C, middle panel) Trichrome and (C, lower panel) Hematoxylin and Eosin (H&E) staining of POD16 tissue used for blinded A and B rejection scoring (bar graphs). The histology shown is representative of at least six transplants per group. (D) Gating strategy to identify CCSP^+^CD326^+^ club cells. Data displayed is a representative result of N=5/group. (E) Intragraft total and indicated subset CD4^+^ and CD8^+^ T cell numbers at POD 16. The bar graph and dot plots show mean ± standard deviation for (C) Mann-Whitney U test and (D, E) Welch’s t-test, where *p <0.05 and ***p <0.001.

Following the induction of lung allograft acceptance, recipients ingested doxycycline on post-operative day (POD) 7 for 18 h and were examined for signs of CLAD development. On POD 16, we observed more histological evidence of peribronchiolar fibrosis in Crry-deficient recipients (**Figure 3C**). Grading for lung transplant rejection using International Society for Heart and Lung Transplantation (ISHLT) criteria in a blinded fashion (44) showed that, compared to lungs engrafted into wildtype mice, allografts transplanted into Crry-deficient recipients had higher “B” airway inflammation. Consistent with these findings, club cells failed to fully reconstitute the graft epithelium in Crry-deficient hosts, with some club cells observed scattered throughout the interstitium, suggesting that epithelial repair is inhibited by augmented complement activation **(Figure 3D)**. Similar to previous findings in humans with CLAD (45, 46), there were moderately higher intragraft T_h_17 and IFN-γ^+^ CD8^+^ T cells in mouse lung recipients lacking Crry (**Figure 3E**).

### Impaired complement regulation promotes intragraft B cell activation and DSA generation

De novo DSA is associated with an increased risk for CLAD development and progression (47). Previous work has shown that activated C3 fragments bound to B cells enhance their activation (48, 49). Given these observations, we asked whether Crry deficiency was associated with increased B cell activation in our CLAD model. C3d deposition on B cells was more prevalent in the spleen and allografts of Crry-deficient recipients (**Figure 4A**). There were also more total and proliferating CD19^+^B220^+^ B cells in the allografts of Crry-deficient recipients (**Figure 4B**). We also quantitated B cell levels in human lung transplant recipients carrying the C3 R102G variant (**Figure 4C**). A sub-cohort of 218 recipients with C3 R102G had increased CD19^+^ B cell frequencies in their bronchoalveolar lavage fluid after adjusting for age, sex, diagnostic group, and ethnicity.

**Figure 4:**
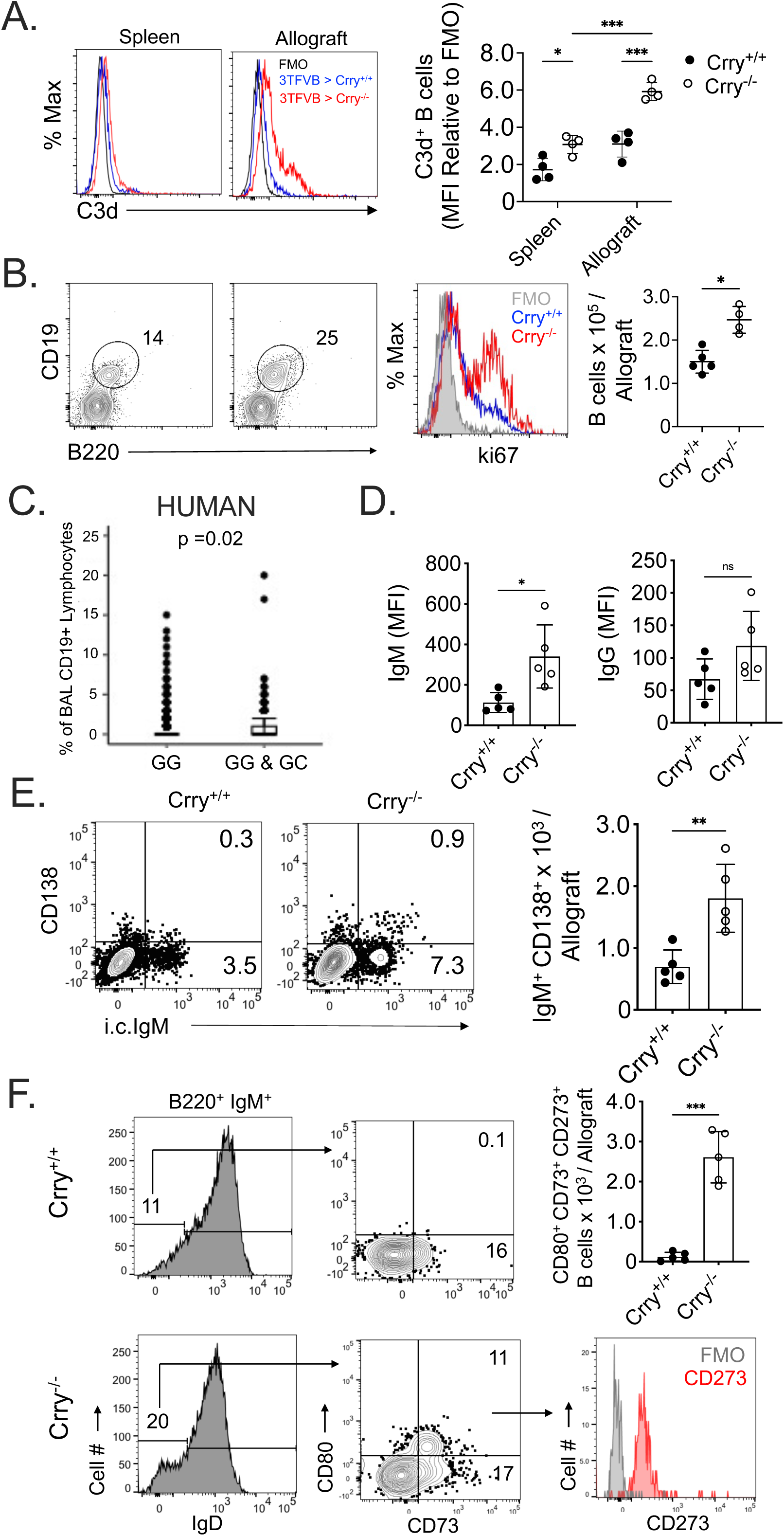
Enhanced intragraft B cell accumulation and activation in lung recipients with a defect in complement regulation. (A) Levels of C3d bound to B cells two days after the induction of doxycycline-induced allograft club cell injury. Representative C3d B cell staining and mean fluorescence intensity plots (N=4/group) using flow cytometry (FACS). (B) FACS analysis of intragraft B cell abundance and ki67 staining (proliferation) with a plot of total B cell numbers on POD 16. FACS contour plots and histograms are representative results from N = 4/group. (C) % of BAL CD19^+^ lymphocytes in UCSF cohorts who carry the GG genotype and C3 R102G (CC&GC) polymorphisms. P=0.02 by an unpaired t-test. (D) Indicated IgM and IgG DSA levels on POD 16 (N=5/group). (E) Intragraft CD138^+^ IgM^+^ antibody-secreting cells abundance as shown by representative FACS dot blots (N=5/group) and bar graphs on POD16. IgM expression was measured by intracellular (i.c.) staining. (F). A representative gating strategy was used to identify intragraft CD80^+^CD73^+^CD273^+^ memory B cells with bar graphs showing the total intragraft numbers of these cells on POD16. Bar graphs and dot blots show means ± standard deviations where (A) 2-way ANOVA with Sidak’s multiple comparisons test, and (B, D, E) Welch’s t-test was conducted where ns is not significant, *p < 0.05, **p < 0.01, *** p < 0.001.

Further analysis of Crry-deficient recipients revealed increased circulating IgM^+^ DSA with a non-significant trend towards elevated IgG^+^ DSA (**Figure 4D**). Recent work has demonstrated that human and mouse lung recipients with CLAD accumulate plasma cells in their graft tissue, implicating a role in local DSA production (14). In line with a higher generation of IgM^+^ DSA, Crry-deficient recipients had more intragraft IgM^+^ antibody-secreting cells than wildtype recipients (**Figure 4E**). CD73, CD86, and CD273 co-expression defines a memory B cell subset poised to become antibody-secreting cells (50). We detected CD73^+^CD80^+^CD273^+^ memory phenotype B cells in allografts after transplantation into Crry-deficient, but not wildtype recipients (**Figure 4F**).

## DISCUSSION

CLAD is characterized by a persistent and irreversible decline in lung function, and is the major cause of long-term morbidity and mortality after lung transplantation (2, 51). Both innate and adaptive immune responses contribute to the progression of CLAD (52). The complement system is an early component of the innate immune response, and has been shown to influence outcomes in primary graft dysfunction, a form of acute lung injury occurring early after lung transplantation (53–55). However, complement activation also influences long-term adaptive immune responses in multiple organ systems and has been implicated in CLAD pathogenesis (9). Complement activation has been shown to augment alloimmune CD4^+^ and CD8^+^ T cell responses through C3a-C3aR interactions (20, 56, 57). However, more specifically, IL-17 activation in lung allografts suppresses membrane regulatory protein expression in airway epithelial cells, and results in increased local complement activation (9). Increased C3a levels enhances IL-17 production, setting up a feedforward loop that worsens obliterative bronchiolitis (OB). Increased C3a levels also upregulate TGF-β, a key mediator of OB, which downregulates membrane regulatory proteins such as CD46/Crry and CD55 in models of lung fibrosis (22, 58). This TGF-β-mediated loss of regulatory proteins is abrogated by both C3aR and C5aR1 inhibition, demonstrating how complement activation is a key component of amplification loops propagating irreversible airway and parenchymal fibrosis.

However, there is considerable heterogeneity in the onset of CLAD. Risk factors such as infection, air pollution, aspiration and a prior history of primary graft dysfunction (PGD) or acute cellular rejection predispose to an earlier onset of CLAD (1, 6, 59). Given the heterogeneity of immune responses, there is an increasing acknowledgment that particular recipients may be genetically predisposed to worse CLAD-free survival (60, 61). The C3 R102G functional polymorphism has been associated with worse outcomes in kidney and liver transplantation (62, 63). Lung transplant recipients who had a donor with C3 R102G demonstrated worse bronchiolitis-obliterans syndrome (BOS)-free survival (64). However, at least in renal transplantation, there has been no benefit to matching donors with recipients based on this polymorphism (65), and long-term lung transplant outcomes in the recipients with advanced lung disease and C3 R102G had not been investigated to date. Surprisingly, we observed nearly 1 in 3 lung transplant recipients harboring the C3 R102G polymorphism, which exceeds the minor allele frequency of 10-15% in the general population, suggesting it also plays a role in the pathophysiology of underlying lung disease. However, having such a common polymorphism among recipients also necessitates a better understanding of how it worsens CLAD-free survival in the setting of impaired complement regulation. Moreover, the prevalence of this functional polymorphism affords avenues to test whether enriching clinical trials in lung transplantation may facilitate personalized immunomodulation to improve CLAD-free survival.

To further probe the effects of dysregulated complement activation on CLAD development, we analyzed graft injury and inflammation in a mouse orthotopic lung transplant model utilizing Crry-deficient mice as recipients. Crry is a membrane complement regulator expressed on a wide variety of murine cells that prevents mouse C3 fragment deposition in response to classical or alternative complement pathway activation (66). Like CD46, Crry regulates complement activation through Factor I-mediated cofactor activity (39, 41). In contrast to wildtype recipients, *Crry^-/-^* recipients had considerably more evidence of graft airway injury and higher levels of DSA, suggesting that CLAD results from antibody-mediated complement activation. However, whether DSA-mediated complement activation is required for graft epithelial injury remains controversial. Introducing either complement-activating or non-complement-activating IgG against donor antigens into a mouse heterotopic tracheal transplant model has been shown to generate obliterative airway disease (67). In the mouse orthotopic lung transplant model, DSA generation occurred de novo approximately eight days after club cell injury. Given the rapid development of CLAD in this model, DSA differences were only evident in the circulating IgM compartment. Compared to IgG, IgM is more efficient at activating the classical pathway due to an approximately thousand-fold higher binding affinity for C1q (68). We have previously demonstrated that club cells are critical to mediating bronchial epithelial repair in mouse lung allograft recipients (43). Club cells also express MHC I and II (43, 69). Therefore, our data suggests that high DSA levels in Crry-deficient recipients could inhibit bronchial repair through complement-mediated injury of club cells.

In addition to causing tissue injury, impaired complement regulation may directly enhance humoral immunity. The complement receptor type 2 (CR2/CD21) is a co-receptor for antigen-bound C3 fragments (48, 70). In mice, co-clustering CR2 with the B cell receptor lowers the threshold of B cell activation, leading to enhanced antibody production (71). CR2 is also expressed on stromal and follicular dendritic cells, where it plays a vital role in antigen retention to promote the generation of B cell memory and antibody-secreting cells (72). However, concerning these reports, the degree to which germinal centers contribute to IgM^+^ memory B cell formation is unclear. Earlier work has suggested that IgM^+^ B cell memory generation can occur without CD40 signaling or T cell help (73). This raises the possibility that our observation of rapid intragraft memory IgM^+^ B cell and DSA accumulation in Crry-deficient recipients is independent of germinal center activity. In addition, previous work has shown that complement fragment deposition on B cells promotes their activation (74). Intriguingly, the B cell receptor C1q binding domain has been shown to drive the fixation of C3 fragments to the B cell membrane and to generate optimal antigen-specific responses (49). Regarding these previous findings, we noted high levels of C3d deposition on intragraft *Crry^-/-^* recipients relative to wildtype recipients. This observation was associated with augmented B cell accumulation, proliferative responses, and enhanced memory B cell and antibody-secreting cell levels. Interestingly, a recent single-cell RNA sequencing study revealed the association between intragraft accumulation of plasma cells within human and mouse CLAD allografts (14).

Collectively, data from the mouse orthotopic lung transplant model implicate dysregulated complement activation in stimulating local B cell activation responses that promote CLAD development.

It is interesting to note that CLAD-free survival in the C3 R102G polymorphism setting occurred primarily in those who developed anti-HLA donor-specific antibodies. However, DSA prevalence did not differ among those with and without the polymorphism. From the standpoint of leveraging this polymorphism for patient selection in future clinical trials, mechanistic studies could involve determining whether C3 R102G worsens allograft injury via a C3d-CR2-mediated axis to promote memory B cell responses. Further investigation into probing impaired classical pathway activation on allograft endothelial surfaces could be addressed with a C1-esterase inhibitor, a C3 inhibitor, or a C5 inhibitor. Given that there are FDA-approved anti-complement therapies that target these molecules, suggesting these therapies could evolve into readily available clinical options for complement-mediated injury in lung transplant recipients (15).

Our study has limitations, including an insufficient sample size at both sites to test whether this polymorphism predisposes to antibody-mediated rejection as a mediator of CLAD. Moreover, the mouse model of Crry deficiency is not entirely analogous to the human complement system. We did not measure other non-HLA anti-allograft antibodies in the allograft or circulation. However, our study’s strengths include its large multicenter investigation with concordant findings in human lung transplant recipients. By utilizing an orthtotopic mouse model of lung transplantation with impaired complement regulation in recipients, we investigated the mechanistic basis of how a genetic polymorphism in a central component of the complement cascade increased complement activation and contributed to CLAD.

Thus, our multicenter study shows that complement activation predisposes to CLAD through impaired regulation. Moreover, the mechanistic basis for this predisposition may primarily be B cell-dependent, suggesting that conventional T cell immunosuppression regimens may be less effective. By providing a sizeable subgroup of lung transplant recipients with a polymorphism that predisposes them to impaired complement regulation, we propose a cohort that can form the basis of clinical trials in personalized immunomodulation to improve long-term outcomes after lung transplantation.

## METHODS

### Sex as a biological variable

Our study investigated data from both male and female human participants, and similar findings are reported for both sexes. Our orthotopic lung transplant model used male donor lungs transplanted into male recipients or female donor lungs transplanted to female recipients.

### Study Design, Settings and Participants

We performed a retrospective cohort study and screened 1218 primary lung transplant recipients at Barnes-Jewish Hospital between January 1, 2005, and December 31, 2022, with follow-up through December 31, 2023. We excluded those who did not consent (n=250) and those in whom genotyping had not been performed (n=576, **Figure 1A**).

At the same time, we also included 864 primary lung transplant recipients at University of California-San Francisco through September 30, 2021. We excluded those who did not consent (n=153) and those in whom genotyping had not been performed (n=293, **Figure 1B**).

### Clinical Variables

Clinical management at these two centers has been described in prior studies (11, 60). HLA antibodies were detected using the LABScreen™ Single Antigen assay, and donor-specific antibody testing (DSA) was considered positive if the reported mean fluorescence intensity (MFI) was ≥2000. The time from transplantation to the first detection of DSA post-transplantation was defined as **time to DSA positivity**. A positive bacterial culture from a bronchial wash or BAL was defined as **bacterial isolation**. PGD and acute cellular rejection were defined based on the International Society of Heart and Lung Transplantation (ISHLT) criteria. CLAD was defined as a persistent decline in FEV1 ≤80% for at least 3 weeks without a specific cause. **CLAD-free survival** was defined as the earliest occurrence of either CLAD or death, whichever came first.

### Genotyping

At BJH, salivary specimens from consented subjects were genotyped for the C3 R102G (*rs2230199*) polymorphism, based on the TaqMan assay, as previously described. At UCSF, blood specimens were genotyped using a transplant-targeted Affymetrix gene array designed by the iGeneTRAiN consortium that directly typed the C3 R102G (*rs2230199*) polymorphism (75). At both centers, recipients with CC/CG (minor allele) were compared to those with GG.

### Bronchoalveolar lavage fluid analysis

Specimens included in this study were from bronchoscopies that were conducted as a part of routine clinical management, when patients had an unexplained decline in their lung function and were being evaluated for CLAD. In the UCSF cohort, bronchoscopies were performed for clinical indication or for allograft surveillance at 0.5, 1, 2, 3, 6, 12, 18 and 24 months after transplantation. Bronchoalveolar lavage cell subsets, including B cells, are routinely quantified by clinical cytometry at UCSF.

### Crry-deficient mice

Crry deficiency has previously been reported as being embryonic lethal because breeding *Crry^+/-^* mice together failed to achieve any *Crry ^-/-^* mice (76–78). However, strategic breeding procedures led to the generation of *Crry ^-/-^* mice. In detail, breeding *Crry^+/-^C3^-/-^*mice resulted in *Crry^-/-^C3^-/-^* offspring, indicating that *Crry ^-/-^* mice can survive in the condition of C3 deficiency. We then bred *Crry^+/-^* males against *Crry^-/-^C3^-/-^* females to generate *Crry^-/-^C3^+/-^* mice. These mice only have half capacity of the alternative pathway activity (41), so we predicted that mothers with the genotype of *Crry^-/-^C3^+/-^* will have insufficient alternative pathway activity to attack embryos that had a *Crry^-/-^* genotype. Considering this nature, we bred *Crry^+/-^* males against *Crry^-/-^C3^+/-^* females, leading to the generation of *Crry^-/-^C3^+/+^* pups, which are *Crry^-/-^* but C3-sufficient. Bronchoalveolar lavage (BAL) was performed on these mice by instilling 1 mL of phosphate-buffered saline (PBS) + enzyme inhibitor (Halt Protease and Phosphatase Inhibitor Single-Use Cocktail, Thermo Fisher, catalog 78442), performing centrifugation at 500g for 5 min, followed by aliquoting the supernatant.

### Immunoblot analysis

C3 was detected using immunoblotting of BAL and serum (MP Biomedicals, catalog 55444, 1:3000). 3 µL of BAL supernatant was mixed with 7 µL of PBS, resulting in a total volume of 10 µL. Subsequently, 3.5 µL of 4x reducing buffer (1:10 ratio of beta-mercaptoethanol added to Laemmli buffer (Bio-Rad, catalog 1610747) was added to the diluted samples, bringing the final volume to 13.5 µL. The samples were vortexed for 5 seconds and then centrifuged. Following centrifugation, the samples were incubated in a heating block at 98°C for 6 min. After this heating step, the samples were centrifuged again before loading onto the gel.

For serum samples, 1 µL of each sample was combined with 9 µL of PBS, also resulting in a total volume of 10 µL. Like the BAL samples, 3.5 µL of 4x reducing buffer was added, and the samples were vortexed, centrifuged, heated, and centrifuged again before gel loading. Purified C3 protein was prepared at a final concentration of 7.5 ng/µL by diluting 1 µL of the purified protein in 99 µL of PBS, followed by the addition of 33.5 µL of 4x reducing buffer.

### Orthotopic left lung transplantation

C57BL/6 (B6) mice were purchased from Jackson Laboratory. CCSP rtTA/TetOCre/DT-A on a mixed background mice were gifts from Dr. Jeffrey Whitsett of The Children’s Hospital of Cincinnati. These mice were previously backcrossed to greater than 99% FVB/J background using microsatellite-assisted accelerated backcrossing (MAX-BAX^SM^; Charles River) to generate 3T FVB left lung donors (43). Left lung orthotopic lung transplantation has been previously described (79). To induce allograft acceptance, recipients received i.p. 250 μg of CD40L Abs (clone MR1) on POD 0 and 200 μg of mouse recombinant CTLA4 Ig on POD2 (80). Club cell injury was triggered by DOX ingestion via food (625 mg/kg chow; ENVIGO) and water (2 mg/mL, MilliporeSigma) for 18 h to induce club cell injury.

### Immunohistological staining

Harvested grafts were formaldehyde-fixed, paraffin-embedded, and stained with H&E or Masson’s trichrome stain. Lung transplant histology was graded by a blinded pathologist using the 2007 revision of the 1996 working formulation for the standardization of nomenclature in the diagnosis of lung rejection (44). For immunohistochemical analysis, paraffin sections were first blocked with 5% goat serum and 2% fish gelatin (both from Sigma-Aldrich) at 25°C for 45 min. Sections were then stained with 1:500 polyclonal rabbit anti-mouse/rat CCSP (catalog WRAB-3950, Seven Hills Bioreagents) and mouse anti-acetylated tubulin, 1:5,000 (clone 6-11B-1, Sigma-Aldrich) overnight at 4°C. For secondary Ab-mediated immunofluorescent visualization, we used 1:1,000 goat anti-mouse Alexa Fluor 488–labeled secondary Abs (catalog A-11-001, ThermoFisher), 1:1,000 donkey anti-goat Alexa Fluor 488 (catalog A-11055, ThermoFisher), 1:1,000 donkey anti-rabbit Alexa Fluor 555 (catalog A-31572, ThermoFisher), and 1:1,000 goat anti-rabbit Alexa Fluor 555 (catalog 4413S, Cell Signaling Technology).

### Flow cytometric analysis

Lung tissue was minced and digested in an RPMI 1640 solution with type 2 collagenase (0.5 mg/mL) (Worthington Biochemical) and 5 units/mL DNAse (MilliporeSigma) for 90 min at 37°C and then filtered through a 70 μm cell strainer (Thermo Fisher) and treated with ACK lysing buffer (Worthington Biochemical). Live cell discrimination was conducted with Zombie (BioLegend) Fixable Dye. Cell surface staining was conducted with the following Abs: CD45 (clone 30-F11; eBioscience), CD45.2 (clone 104; BioLegend), CD90.2 (clone 53-2.1; eBioscience), CD4 (clone RM4-5; eBioscience), CD8α (clone 53-6.7; eBioscience), B220 (RA3-6B2; Biolegend), CD19 (1D3; BD Biosciences), Goat-anti mouse/rat C3d (Bio-Techne), CD138 (281-2; Biolegend) CD38 (S21016F; Biolegend), IgD (11-26c.2a; Biolegend), Foxp3 (FJK-16s; Thermofisher), CD73 (TY/11.8; Biolegend), CD80 (16-10A1; Biolegend), CD273 (TY25; Thermofisher), CD31 (clone 390; BioLegend), CD34 (clone HM34; BioLegend), and CD326 (clone G8.8; BioLegend). Intracellular CCSP (Seven Hills Bioreagents) and polyclonal goat anti-mouse-IgM-heavy chain (SouthernBiotech) staining was conducted with the Cytofix/Cytoperm kit (BD Biosciences) following the manufacturer’s recommendations. Staining for Foxp3 (FJK-16s, eBioscience), Ki-67 (16A8; BioLegend), was conducted with the Intranuclear Transcription Factor Staining Buffer Kit (Invitrogen) per the manufacturer’s recommendations. For IFN-γ and IL-17A expression, cells were first stimulated with 1 μM ionomycin (MilliporeSigma) and 20 ng/mL PMA (MilliporeSigma) for 3.5 h, with 2 μM Golgi Plug (BD Biosciences) added for the last 3 h of stimulation. Cells were then intracellular stained with IFN-γ (clone XMG1.2; eBioscience) and IL-17A (clone TC11-18H10.1; BioLegend) using a Cytofix/Cytoperm kit. For DSA analysis, FVB splenocytes were resuspended at 4 x 10^6^ cells/mL in FACS buffer (PBS, 0.1% BSA,0.02% NaN_3_) and treated with FcγR block on ice for 10 min (93; Biolegend). Serum from transplanted mice was diluted to 1/25 in PBS, and 50 μL of this diluted serum was mixed with 50 μL of FVB splenocytes, which were then incubated for one hour at 4^0^C. Cells were washed 2x with FACS buffer and stained with Goat anti-mouse-IgM-heavy chain-APC or Goat anti-mouse-IgG-Fcγ-fragment-AlexaFluor ^488^ (Jackson ImmunoResearch), and washed twice more before FACS and subsequent MFI analysis.

### Statistics

Data were analyzed by the Mann-Whitney U test or 1-way ANOVA and are represented as mean ± SD. Statistical analysis was conducted in R (v4.3.2) and GraphPad Prism software, version 9.0. P < 0.05 was considered significant. Time to event analyses were performed using Kaplan-Meier curves with log-rank test for equality. Multivariable analyses were conducted via Cox Proportional Hazards Regression.

### Study approval

Animal experiments were conducted in accordance with an approved IACUC protocol (Washington University, 19-0827). The Barnes-Jewish Hospital (BJH) participants provided written informed consent in accordance with the Washington University School of Medicine Institutional Review Board for Human Studies protocol (ID # 201811073, 201105421). The University of California-San Francisco (UCSF) participants provided written informed consent in accordance with the Institutional Review Board for Human Studies protocol (ID # 13-10738).

## Data availability

The deidentified data will be made publicly available at the time of publication. The study participants have not consented to sharing genetic data publicly and hence, this cannot be uploaded to a publicly accessible server.

## AUTHOR CONTRIBUTIONS

Designing research studies: H.S.K., D.R.C., J.R.G., A.E.G.

Conducting experiments: H.S.K., F.L., Z.L., L.G., D.H.K., X.W., A.E.G.

Acquiring data: H.S.K., L.K.T., D.R.C., F.L., Z.L., L.G., N.S., C.C., H.J.H., C.A.W., R.R.H.

Analyzing data: H.S.K., L.K.T., D.R.C., C.B.,

Providing reagents: X.W., D.H.K., D.E.B., D.K., J.P.A.

Writing the manuscript: H.S.K., L.K.T., D.R.C., L.G., X.W., J.R.G., A.E.G.

Funding acquisition: H.S.K., L.K.T., D.R.C., D.K., J.P.A., J.R.G., A.E.G.

## ACKNOWLEDGEMENTS

The authors thank Dorit Daphna-Iken, Jihong Zhu, Rafael Aponte Alburquerque and Ayse Naz Ozanturk for their technical assistance.

## Sources of support

**H.S.K.:** National Institutes of Health (R01-HL166449, R01-HL169860), Children’s Discovery Institute, Longer Life Foundation

**L.K.T.:** National Institutes of Health (K01-HL155231), Robert Wood Johnson Foundation, Doris Duke Charitable Foundation

**D.R.C.:** VA Office of Research and Development (BX005301), Cystic Fibrosis Foundation (CALABR22G0)

**J.P.A.:** National Institutes of Health (R35-GM136352)

**J.R.G.:** National Institutes of Health (R01-HL161048, R01-HL151552), Cystic Fibrosis Foundation (GREENL21AB0), VA Office of Research and Development (CX002011)

**A.E.G:** National Institutes of Health (P01-AI116501, R01-HL167277), Barnes-Jewish Foundation.

